# LeafByte: A mobile application that measures leaf area and herbivory quickly and accurately

**DOI:** 10.1101/777516

**Authors:** Zoe L. Getman-Pickering, Adam T. Campbell, Nicholas Aflitto, Todd A. Ugine, Ari Grele, Julie Davis

## Abstract

1. In both basic and applied studies, quantification of herbivory on foliage is a key metric in characterizing plant-herbivore interactions, which underpin many ecological, evolutionary, and agricultural processes. Current methods of quantifying herbivory are slow or inaccurate. We present LeafByte, a free iOS application for measuring leaf area and herbivory. LeafByte can save data automatically, read and record barcodes, handle both light and dark colored plant tissue, and be used non-destructively.
2. We evaluate its accuracy and efficiency relative to existing herbivory assessment tools.
3. LeafByte has the same accuracy as ImageJ, the field standard, but is 50% faster. Other tools, such as BioLeaf and grid quantification, are quick and accurate, but limited in the information they can provide. Visual estimation is quickest, but it only provides a coarse measure of leaf damage and tends to overestimate herbivory.
4. LeafByte is a quick and accurate means of measuring leaf area and herbivory, making it a useful tool for research in fields such as ecology, entomology, agronomy, and plant science.

## Introduction

The amount of leaf tissue consumed, hereafter “herbivory”, is a fundamental metric used to understand plant-herbivore interactions in many disciplines spanning basic and applied science, including plant chemistry, plant-insect ecological and evolutionary dynamics, plant breeding, agronomy, and horticulture (Turcotte et al 2014). However, efficiently and accurately measuring amounts of herbivory remains challenging (Williams et al. 1991).

Herbivory from chewing insects is measured with software such as ImageJ (Abràmoff et al. 2004), mobile apps such as BioLeaf (Machado et al., 2016), and manual methods such as grid quantification (Coley 1983) or visual estimation (Johnson et al. 2016). While all of these methods have advantages, there is significant room for improvement. One of the most commonly used options, the image processing program ImageJ, is accurate but not optimized for measuring herbivory, and is therefore incredibly time-consuming. Further, images must be scanned or photographed, saved on a computer, and then uploaded, which is also slow. The mobile app BioLeaf (Machado et al., 2016) allows for quick and efficient measurements of herbivory. However, it only measures percent herbivory, and not the absolute leaf area and herbivory, making it difficult to compare levels of herbivory when leaf sizes vary, which is commonly the case. Grid quantification entails placing a grid under a damaged leaf and counting the number of squares where an herbivore removed leaf tissue (Coley 1983). While measuring small amounts of herbivory is straightforward, measuring large amounts of herbivory or leaf area can be prohibitively slow. Finally, visual estimation of herbivory is quicker but sacrifices accuracy (Johnson et al. 2016).

We introduce LeafByte, a free and open source mobile app that solves common issues with the current tools and provides additional features. LeafByte can scan barcodes, measure light colored petals or leaves, and save results (with the date, time, and GPS coordinates) to a spreadsheet on the phone or on Google Drive. LeafByte can be used non-destructively. We present a systematic comparison of the accuracy and efficiency of LeafByte and four of the most common herbivory measurement tools: ImageJ, BioLeaf, grid quantification, and visual quantification.

## Methods

### How LeafByte works

Users take or upload an image of a leaf surrounded by 4 dots in a square that act as a scale (see Supporting Information 1). LeafByte identifies the leaf and scale markings by separating the foreground of the image from the background in a process called “thresholding” (Otsu, 1979). Each pixel in the image is considered individually. If the luma of the pixel’s color, a measure of perceived intensity (ITU-R, 1982-2015), is above a certain cutoff value (the “threshold”), that pixel will be considered foreground; otherwise, it becomes background. Because the leaf and scale markings are much darker than the background (typically a green leaf and black scale markings on white paper), they are marked as foreground, while the rest is marked as background. LeafByte also supports light tissue (such as white flowers) against dark backgrounds by simply reversing the process.

LeafByte determines the luma level that separates foreground from background using an algorithm called Otsu’s method (Otsu, 1979). Otsu’s method considers a histogram of lumas in the image. This histogram is typically bimodal, with a mode of high luma, representing the leaf and scale markings, and a mode of low luma, representing the background. Otsu’s method finds a luma that most clearly separates those two modes, effectively distinguishing foreground from background. This automatically-determined threshold is generally effective, but LeafByte allows users to tweak as needed (Fig. 1A).

Next, LeafByte determines what pixels represent the leaf and scale markings using an algorithm called connected-component labeling (Rosenfeld & Pfaltz, 1966) to separate pixels into groups representing different objects. LeafByte assumes that the largest group is the leaf, and the next four largest are the scale markings. This is right in most cases, and when it is not (e.g. there is another object in the image), the user can correct LeafByte’s assumption by manually identifying scale markings (Fig. 1B).

If the image was taken at an angle, the scale markings no longer form a square, and the leaf itself is distorted, causing error (Supporting Information 2). To correct this skew, LeafByte uses a technique called planar homography (Wang, Klette, & Rosenhahn, 2006) to re-distort the image so that the scale markings once again form a square. LeafByte uses connected-components labeling again on background pixels to identify the holes within the leaf.

The user can draw missing margins onto the leaf image (Fig. 1C). Then, counting the number of pixels in the leaf and in the holes gives the relative amount of leaf eaten. Summing the number of pixels in the leaf and the holes gives the total size of the original leaf in pixels. Because there is a known distance between each scale mark, LeafByte can convert numbers of pixels into real world units. The photo and results are saved in a CSV file to Google Drive or the phone.

### Methods for Testing LeafByte

#### Accuracy

To confirm the accuracy of ImageJ and LeafByte, we used both methods to measure artificial “leaves” of known area”. We printed out 16 black rectangles of known area with white “holes” of known size and analyzed them with both LeafByte and ImageJ, comparing theand compared their results to the known area.

#### Comparisons of different methods

We collected 67 leaves from 14 plant species (Supporting Information 3) from the Cornell Botanical Garden and grounds. Leaves were selected to represent a range of morphologies and were categorized by shape. If the leaf was undamaged, we created artificial herbivory using hole punches and razor blades to remove 0-50% of the leaf. We recorded whether the leaf was damaged on the margin (n=36) or only internally (n=22). Herbivory was estimated visually and using grid quantification (Coley 1983). For visual estimation, herbivory was estimated to the nearest 5%. Leaves with 0-2.5% herbivory were rounded to 5%. The leaves were then flattened between a sheet of printer paper with the scale printed on it and a Premium Matte Film Shield Screen Protector (J&D, Middleton, MA) and photographed. Each photograph was analyzed using LeafByte, BioLeaf, and ImageJ by at least two different researchers per method. LeafByte and ImageJ provided total leaf area, absolute herbivory, and percent herbivory. BioLeaf and visual quantification provided only percent herbivory, and the grid method provided only percent herbivory. We also recorded the time it took to analyze each leaf and record the data. For ImageJ, we did not include the time it took to photograph and upload the pictures.

### Statistics

All statistics were performed using R, Version 3.5.2 (R Core Team, 2018). We built global mixed effects models using the nlme package (Pinheiro et al., 2018). We dropped non-significant predictors from the models in a backwards stepwise fashion, assessed pairwise differences between the methods using emmeans (Lenth, R., 2019), and adjusted for multiple comparisons using false discovery rate.

#### Accuracy

To test for differences in measurement accuracy between ImageJ and LeafByte, we ran linear mixed effects models with area and herbivory as response variables. In both models, method was included as a fixed effect, and the known size of each artificial leaf was set as the reference value. Additionally, we used an equivalency test (TOSTER, Lakens 2017) to evaluate whether the methods produced the same results (as opposed to linear models that test for differences). We used ¼ of the standard deviation as upper and lower bounds of the model.

#### Comparisons of different methods

To analyze the effect of method on leaf area, we ran a linear mixed effects model with leaf area as the response variable and the interaction between method and leaf shape as predictor variables. Species and leaf ID were included as random effects in all models. Leaf areas were log transformed to meet assumptions of homoscedasticity.

To analyze the effect of method on herbivory, we ran a linear mixed effects model with herbivory as the response variable and the interaction between method and number of holes and the interaction between method and presence of leaf margin herbivory as predictor variables. To analyze the effect of method on percent area consumed data, we ran a binomial generalized linear mixed effects model with herbivory as a response variable and the interaction between method and number of holes and the interaction between method and presence of leaf margin herbivory as fixed effects. Because low levels of herbivory (0-2.5%) were rounded to 5% rather than 0% when using visual quantification, we analyzed both the full data set and data where percent herbivory was greater than 5% to ensure that rounding did not skew our results.

## Results

### Accuracy

We found no difference between the known area and LeafByte for total area (t-ratio=0.126, df=36, p=0.991, Fig. 2A) or herbivory (t-ratio=1.11, df=36, p=0.512, Fig. 2B) or between the known area and ImageJ for total area (t-ratio=-1.53, df=36, p=0.285, Fig. 2C) or herbivory (t-ratio=0.793, df=36, p=0.710, Fig. 2D). On average, LeafByte differed from the known area by 1.3% while ImageJ differed from the known area by 3.2%. Based on the equivalence test comparing LeafByte to the known area, we can conclude that the difference between the treatments is equivalent to zero (t_36_=20.4, p<0.001, t_36_=-4.40, p<0.001) for both leaf area and hole area. Similarly, the difference between ImageJ and the known area is equivalent to zero for both leaf area and hole area (t_36_=-20.2, p<0.001, t_36_=-4.52, p<0.001).

### Comparisons of different methods

On average, leaf area measured by LeafByte was 2% lower than the leaf area measured by ImageJ (t_248_=0.627, p=0.023, Fig. 3A). There was no effect of leaf shape on leaf area measurements using LeafByte or ImageJ (log likelihood=221 on 8 df, p=0.565). There was a significant interaction between method and number of holes in a leaf on the area of herbivory measurements (log likelihood = 979 on 8 df, p=0.003), such that herbivory was underestimated when there were more holes using the grid method (t_322_=-3.34, *p*=0.001), but not any of the other methods. When holding hole number constant, there was no significant difference in herbivory estimates between ImageJ and LeafByte (t-ratio=0.002, df= 322, *p*=1.0) or ImageJ and grid quantification (t-ratio=-2.02, df= 322, p=0.110, Fig. 3B). There was a significant effect of method on percent herbivory (F_3,107_= 35.8 p<0.001, Fig. 3C). Neither BioLeaf (z-ratio=-0.871, p=0.820) nor LeafByte (z-ratio= -0.955, p=0.775) were significantly different from ImageJ. Visual quantification overestimated percent herbivory compared to ImageJ (z-ratio= -5.12, p<0.001), LeafByte (z-ratio=4.87, p<0.001), or BioLeaf (z-ratio=-4.867, p<0.001). The accuracy of each method was not affected by the presence of margin herbivory (log likelihood= -767 on 14 df, *p*=0.102) or the number of holes (log likelihood = -770 on 10 df, *p*=0.912). The results were the same when analyzing the full data set or only the data >5%.

Different methods took different amounts of time to analyze a given leaf (F_4,549_=202, p<0.001, Fig. 3D). ImageJ was by far the slowest option, taking twice as long as LeafByte (t-ratio=-15.0, df=549, p<0.001) on average. Grid quantification and LeafByte took a comparable length of time (t-ratio=-0.508, df=549, p=0.612). BioLeaf was 40% faster than LeafByte (t-ratio=5.41, df=549, p<0.001) while visual quantification was 85% faster (t-ratio=11.7, df=546, p<0.001). The presence of margin herbivory slowed down leaf measurements for LeafByte (t-ratio=-3.14, df=52, p=0.003), ImageJ (t-ratio=-3.79, df=52, p<0.001), and BioLeaf (t-ratio=-2.67, df=52, p=0.0010), but not the grid method (t-ratio=-1.69, df=52, p=0.097) or visual quantification (t-ratio=0.655, df=52, p=0.515). The number of holes increased the time to analyze for all methods (F_4,549_=10.0, p<0.001), although it was drastically higher for ImageJ, which took ∼8 seconds per additional hole, while all other methods were less than ½ a second per hole.

## Discussion

LeafByte is a novel tool that combines and improves on the strengths of existing tools in a user-friendly application. LeafByte quickly and accurately measures leaf area, herbivory from chewing herbivores, and percent herbivory. It is the first herbivory measurement app to automatically save measurements to a spreadsheet, reducing time and transcription errors. LeafByte can read and record barcodes, handle both light and dark colored plant tissue, and be used non-destructively. Our testing illustrates that while LeafByte produced average measurements 2% lower than ImageJ, both LeafByte and ImageJ were highly accurate when measuring “leaves” and “herbivory” of known sizes. LeafByte takes half as long as ImageJ to measure each leaf and can handle larger numbers of holes much more quickly. All electronic methods were significantly slower with margin damage.

We found that visual quantification led to overestimations. This was likely due to lack of training and the fact that most of our leaves had low levels of herbivory (Johnson et al. 2016). Tilting a phone/camera more than 15° caused high rates of error. Using a skew-correcting box as a scale rather than a line was an effective and necessary means of reducing error (Supporting Information 2). Researchers using digital methods that do not automatically correct for skew should take care to ensure that their photographs are not taken at an angle greater than 15%. LeafByte has several limitations. It is difficult to identify margin damage on needles and highly complex leaves. Highly ruffled or complex leaves have more shadows and are difficult to lie flat without overlap. Poor quality photos or photos with extensive shadows make it difficult to cleanly remove the background. These limitations hold for other image processing software including ImageJ and BioLeaf.

While LeafByte was designed to measure leaf area and herbivory, it can also measure disparate things like damage on butterfly wings, fungal growth on petri dishes, insect droppings on filter paper, and the efficacy of anilox rollers. LeafByte is a quick and accurate means of measuring leaf area and herbivory, making it a transformative tool for a wide variety of applications.

## Acknowledgments

We would like to thank Dr. Heather Grab, George Stack, Jose Rangel, Fiona MacNeil, and Danielle Rutkowski for giving feedback on LeafByte; Angeliki Cintron, Abby Grace-Ditmar, Tait Stevenson, Emma Weissburg, and Christina Zhao for collecting data for this paper; Christina Zhao for graph aesthetics; Dr. Eric Raboin for projective geometry help; and Dr. Heather Grab for statistical advice and editing. Authors have no conflict of interest.

## Author Contributions

ZGP and AC designed the and created the app. ZGP, NA, TU, AG, and JD contributed to testing and improving the app. ZGP and JD collected and analyzed the data. ZGP, AC, NA, TU, JD, AG contributed to writing and editing the paper.

---

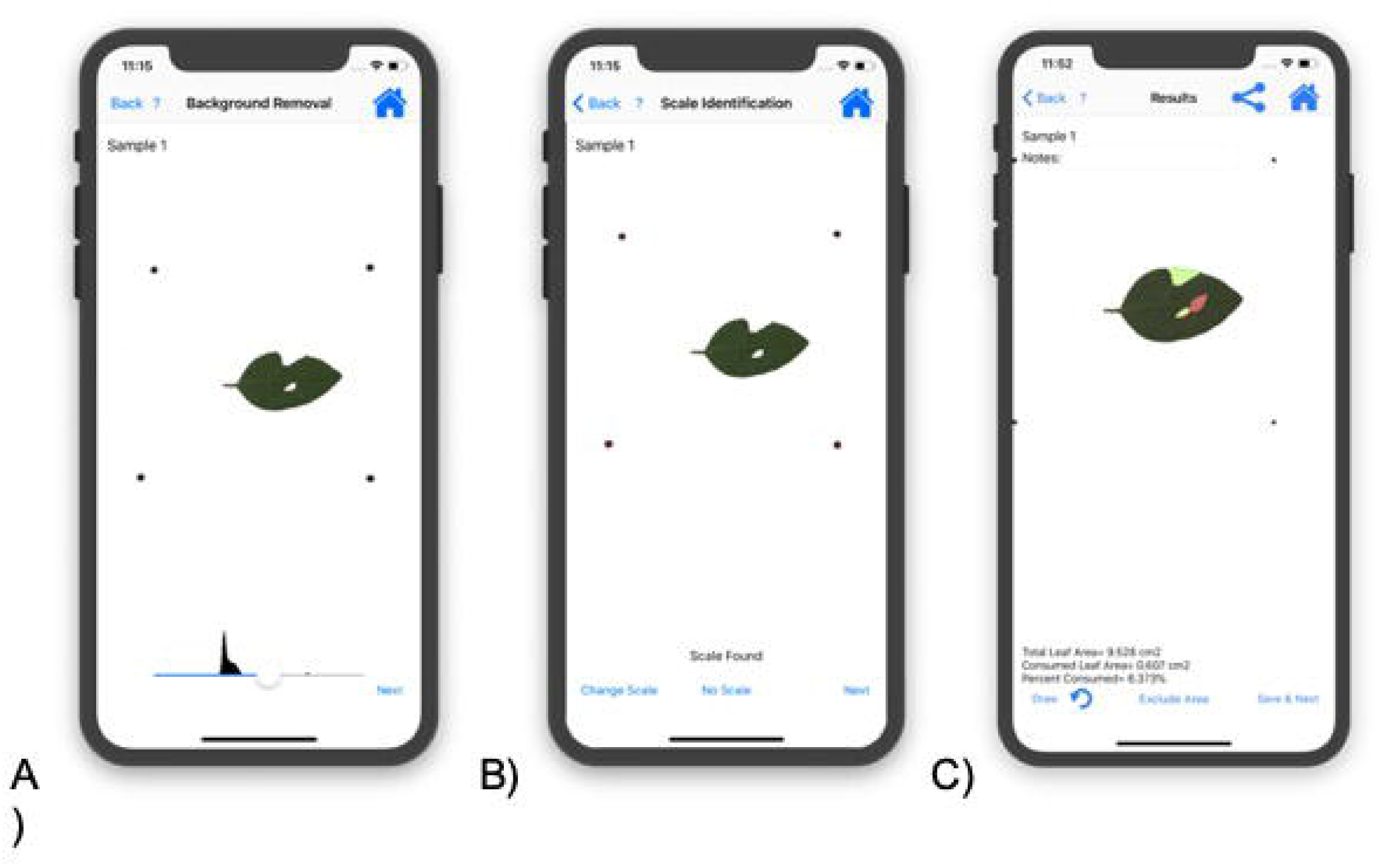

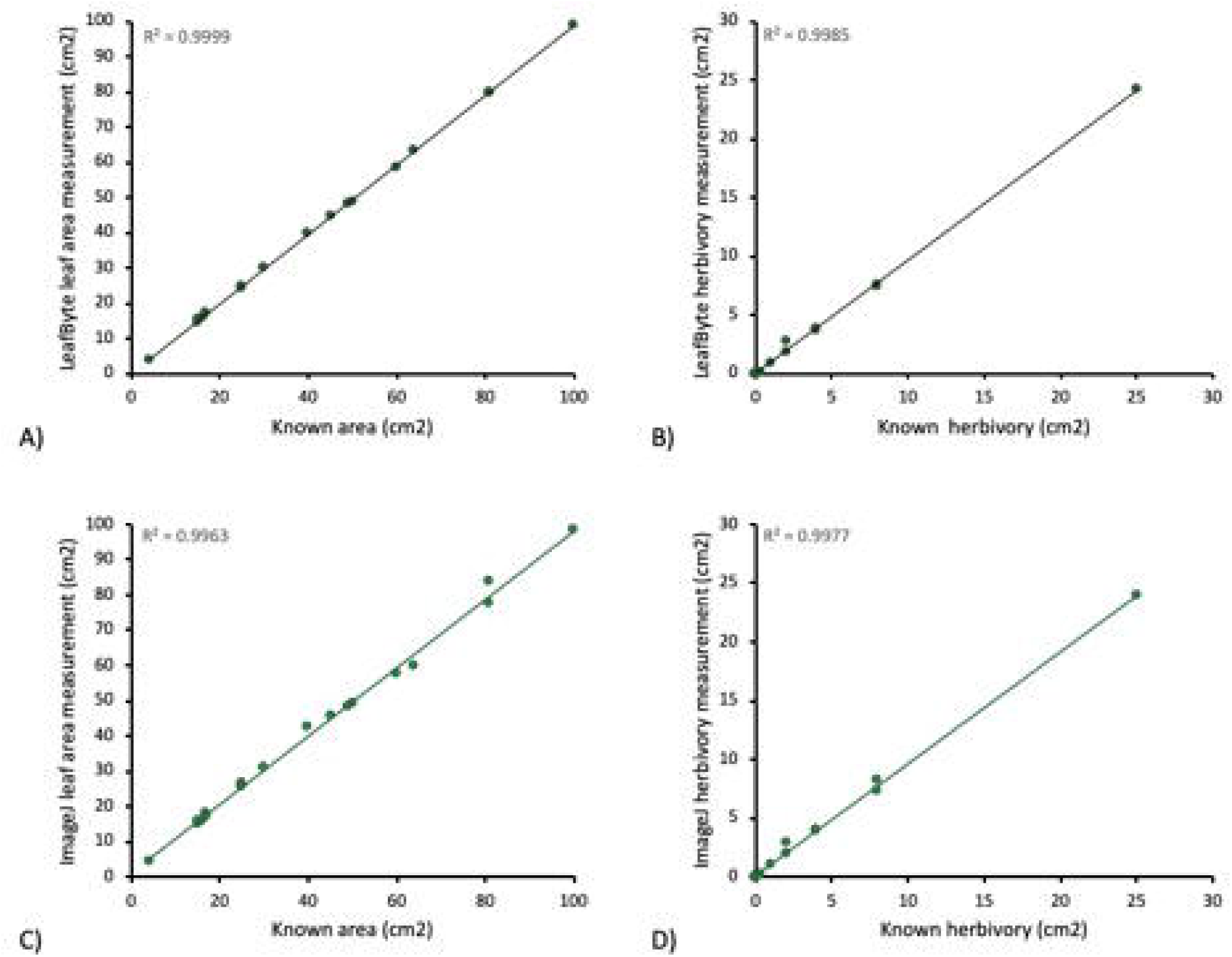

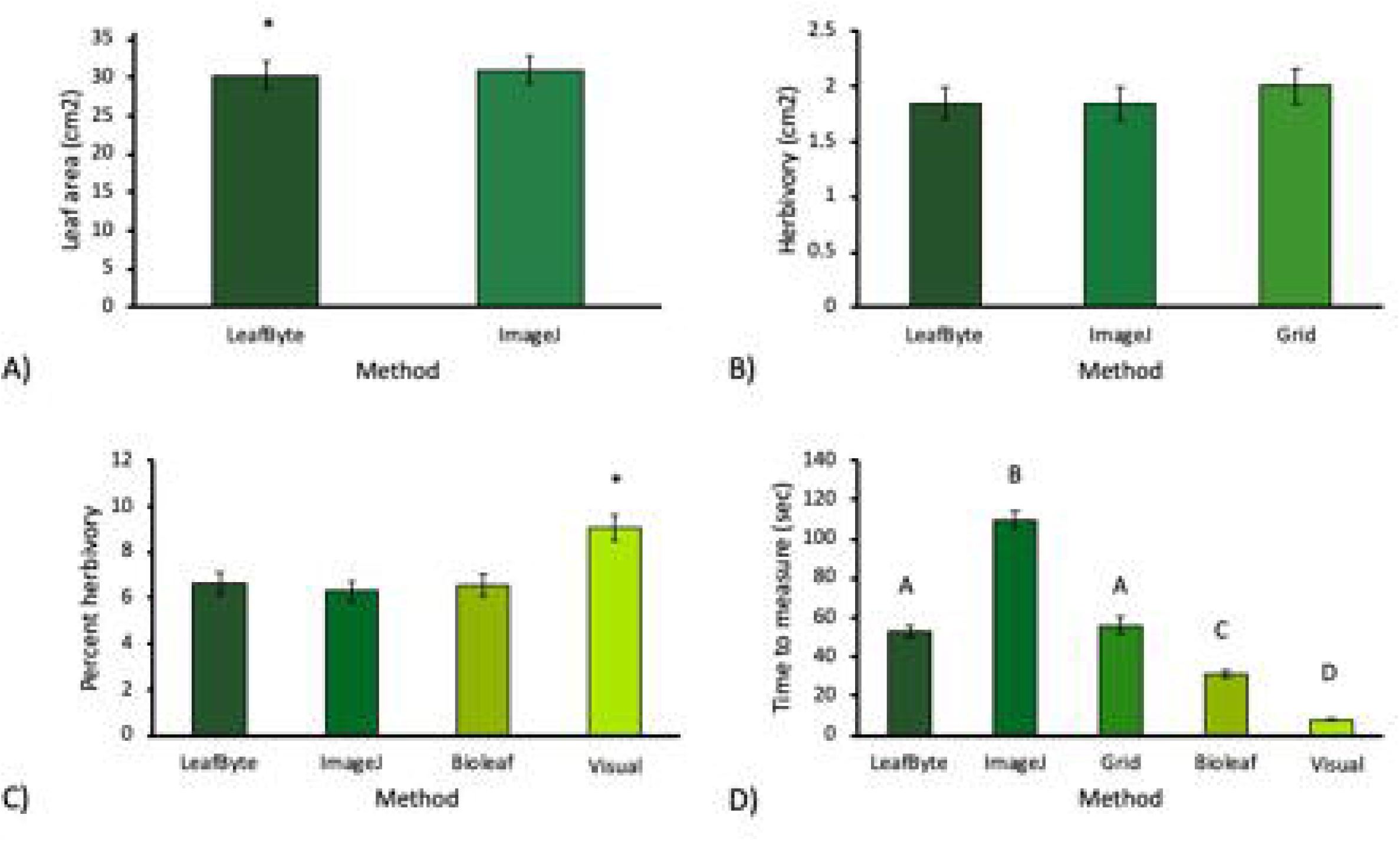

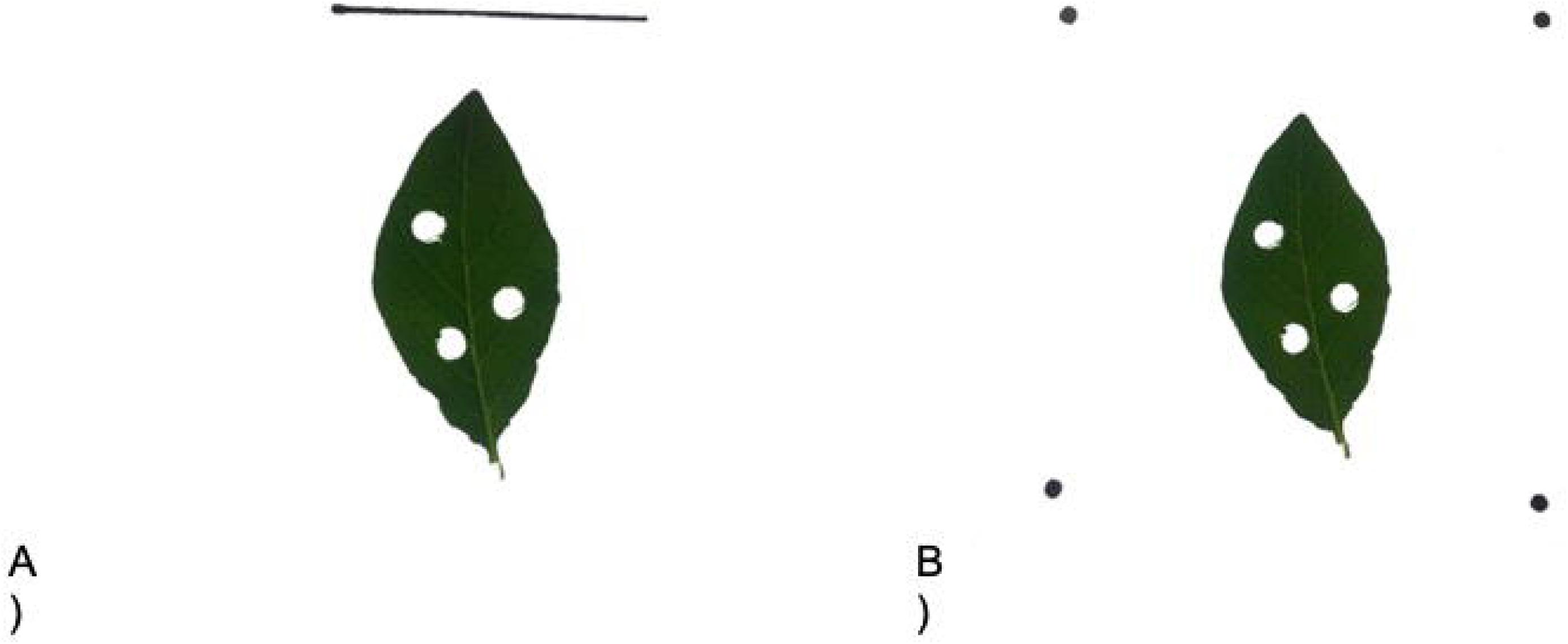

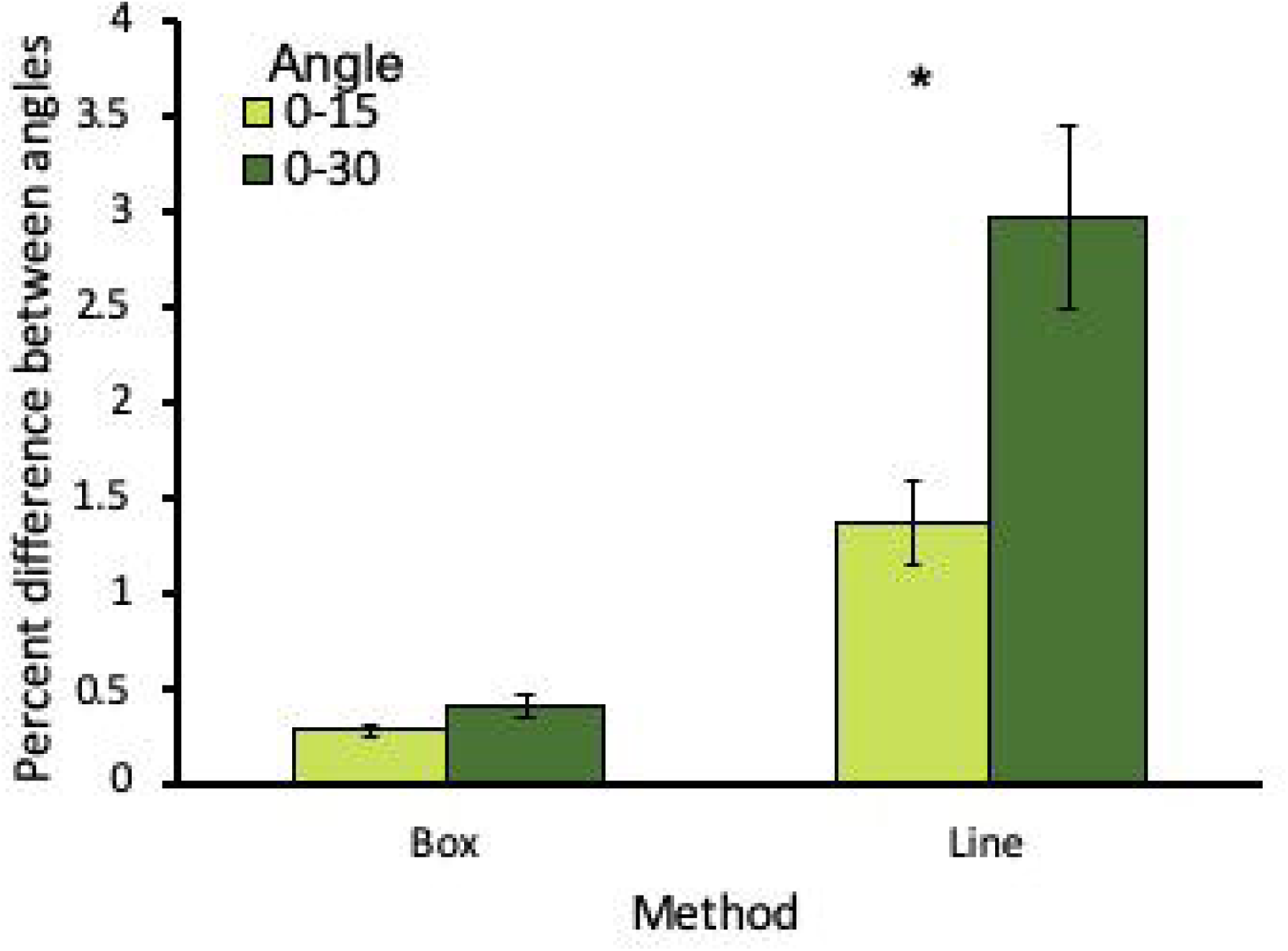

## References

Abràmoff, M. D., Magalhães, P. J., & Ram, S. J. (2004). Image processing with ImageJ. Biophotonics international, 11(7), 36–42.

Coley, P. (1983). Herbivory and defensive characteristics of tree species in a lowland tropical forest. Ecological Monographs, 53(2), 209–229.

International Telecommunication Union Radiocommunication Sector [ITU-R]. (1982-2015).

Recommendation ITU-R BT.601-7. Accessed August 19, 2019. https://www.itu.int/dms_pubrec/itu-r/rec/bt/R-REC-BT.601-5-199510-S!!PDF-E.pdf

Johnson, M. T., Bertrand, J. A., & Turcotte, M. M. (2016). Precision and accuracy in quantifying herbivory. Ecological Entomology, 41(1), 112–121.

Lakens, D. (2017). Equivalence tests: A practical primer for t-tests, correlations, and meta-analyses. Social Psychological and Personality Science, 8(4), 355–362.

Lenth, R. (2019). emmeans: Estimated marginal means, aka least-squares means. R package version 1.3.3. https://CRAN.R-project.org/package=emmeans

Machado, B. B., Orue, J. P. M., Arruda, M. S., Santos, C. V, Sarath, D. S., Goncalves, W. N., … Rodrigues-Jr, J. F. (2016). BioLeaf: A professional mobile application to measure foliar damage caused by insect herbivory. Computers and Electronics in Agriculture, 129, 44–55.

Otsu, N. (1979). A Threshold Selection Method from Gray-Level Histograms. IEEE Transactions on Systems, Man, and Cybernetics, 9(1), 62–66.

Pinheiro J, Bates D, DebRoy S, Sarkar D, R Core Team (2018). _nlme: Linear and Nonlinear Mixed Effects Models_. R package version 3.1-137, <URL:https://CRAN.R-project.org/package=nlme>.

R Core Team (2018). R: A language and environment for statistical computing. R Foundation for Statistical Computing, Vienna, Austria. https://www.R-project.org/.

Rosenfeld, A. L., & Pfaltz, J. L. (1966). Sequential operations in digital picture processing. Journal of the ACM, 13(4), 471–494.

Turcotte, M. M., Thomsen, C. J., Broadhead, G. T., Fine, P. V., Godfrey, R. M., Lamarre, G. P., Meyer, S. T., Richards, L. A. and Johnson, M. T. (2014), Percentage leaf herbivory across vascular plant species. Ecology, 95: 788–788.

Wang, X., Klette, R., & Rosenhahn, B. (2006). Geometric and photometric correction of projected rectangular pictures. Image and Vision Computing.

Williams, M.R. & Abbott, I. (1991) Quantifying average defoliation using leaf-level measurements. Ecology, 72, 1510–1511.

