## Supporting Information 1 for "LeafByte: A mobile application that measures leaf area and herbivory quickly and accurately"

**Supporting Information 1: How to use the app**

*See* [*https://zoegp.science/leafbyte-how-to-use*](https://zoegp.science/leafbyte-how-to-use) *for up-to-date and thorough instructions with pictures.*

Before analyzing leaf area and herbivory using LeafByte, users can customize settings on the settings page. These settings include: where to save herbivory estimates and leaf images, the name of the dataset, the size of the scale, the sample number, the color of the background the photo is taken against (black or white), and whether to scan barcodes. Photos must be taken against a highly contrasting background. Dark plant tissue (e.g. most leaves) should be photographed against a white background, while light plant tissue (e.g. flower petals) should be photographed against a black background. In both cases, the background should include a defined scale comprised of four dots representing the corners of a square (available as a PDF at http://zoegp.science/leafbyte-how-to-use). If absolute herbivory is unneeded, and percent herbivory is sufficient, then a plain background without a scale can be used. Although the scale can be any size, the entire leaf must be contained within the boundary of the square defined by these dots (Fig.1). LeafByte autocorrects skew in images taken at an angle, which is a common user error with handheld cameras. Once a photo has been selected, LeafByte automatically removes the background, letting the user tweak if needed (Fig. 2 A). Next, LeafByte automatically identifies the scale points (indicated by four red dots) and the leaf (indicated by a red leaf symbol) (Fig. 2 B). Users may modify the leaf and scale selection. LeafByte will then automatically measure the total leaf area, consumed leaf area, and percentage of leaf consumed (Fig. 2 C). If herbivory reaches the edge of the leaf, users can zoom in and out and freehand draw margins along the leaf edge. This allows much more flexibility than other methods, especially for leaves with unusual shapes or non-entire margins. LeafByte automatically updates the results.

Areas identified as holes, colored a light green, can be ignored by pressing the “Exclude Area” button and touching the area to be excluded. This is particularly useful when analyzing compound leaves where the leaflets overlap. Notes may be entered via a keyboard at the top of the screen. Once all areas of herbivory are identified, users can press “Save” at the bottom of the screen to save all measurements, notes, and the current image to the selected location (a spreadsheet on the phone or Google Drive). The user may then choose the next photo for analysis, which will be added to the end of the spreadsheet.

***Additional features, and data output:***

LeafByte has additional features configured on the settings page on the home screen:

Data saving - All data can be saved to a CSV file on the phone or as a Sheet in Google Drive. The pictures analyzed by the app can also be saved to the phone or in Google Drive.

Barcode scanning - LeafByte can scan all major 1D and 2D barcodes. The information encoded in the barcodes will be included in any data output files. This feature makes it easier to track individual plants for high throughput herbivory measurements.

GPS - LeafByte can also record the GPS location where the image was processed. LeafByte cannot extract GPS information for images taken previously and then analyzed in LeafByte. It can only measure the GPS location of your phone when the image is processed. LeafByte uses the phone GPS to get a position accurate to within 10 meters. However, the GPS feature can slow saving time on some devices.
