## Supporting Information 2 for "LeafByte: A mobile application that measures leaf area and herbivory quickly and accurately"

**Supporting Information 2: Scale methods**

Initially, we used a straight line of known length as the scale (Fig. 4A), similar to how ImageJ is often used. Testing and observations found that holding a phone at an angle (which happens commonly) leads to skewed results. To fix this problem, we designed a system with 4 dots in a square acting as the scale. This allows LeafByte to identify and correct skew. To compare the line and dots scale methods and validate that our skew-correction resulted in more accurate output, we photographed 46 leaves (Table 1) at 0, 15, and 30 degree angles as determined by the mobile level application ‘Bubble level for iPhone’ (version 3.04). The leaves were photographed and analyzed using LeafByte (version 0.0.7) using a line as a scale, or LeafByte (version 1.0.0) using 4 dots in a square as a scale.

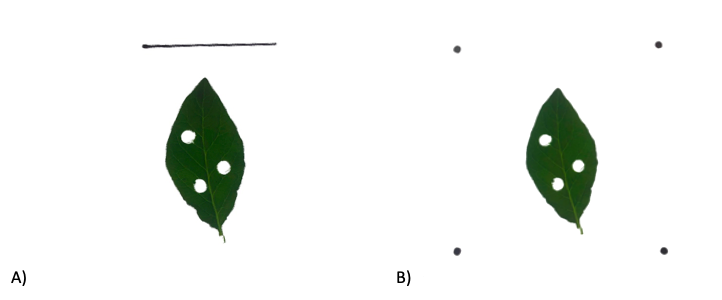

Figure 4. Images of leaves analyzed with A) a line as a scale or B) 4 dots in a square as the scale.

We found a significant scale method by angle interaction on the total herbivory (F_2,199_=10.9, p<0.001). While using a line for the scale, taking the photo at a 30 degree angle altered the herbivory by an average of 13.0% and up to 57.9% (t_,199_= -5.70 p<0.001). Using the dots method, there was no effect of angle between 0-15 (t_199_=-0.118, p=0.942), 15-30 (t_199_=0.327, p=0.943) or 0-30 (t_199_=0.205, p=0.977) degrees on leaf area. There was no effect of angle (F_2,201_=0.760, p=0.469) or scale method (F_1,92_=0.869, p=0.353) on herbivory.

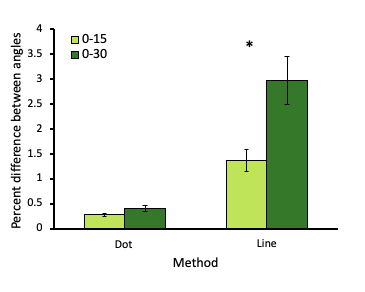

Figure 5. Taking a photo at 30 degree angle significantly skewed the results when using the line as a scale but not when using the dots.

Table 1. Species used in testing the line and dot versions of LeafByte.

| **Code** | **Species** |
| --- | --- |
| CIN | *Potentilla sp.* |
| HR | *Rosa rugosa* |
| GD | *Cornus foemina* |
| WH | *Hamamelis × intermedia* |
| PW | Salix caprea |
| PAP | *Acer griseum* |
| ROD | *Cornus sericea* |
| LP | *Ligularia palmatiloba* |
| SD | *Cornus sp.* |
| PL | *Plantasia* |
| JS | *Spiraea japonica* |
| TE | *Thalictrum* 'Elin' |
| PP | *Parrotia persica* |
| RS | *Rosa sp* |
| HO | *Helleborius odorus* |
| CS | *Cornus sericea* |
| SB | *Scilla bifola* |
