## Supporting Information 3 for "LeafByte: A mobile application that measures leaf area and herbivory quickly and accurately"

**Supporting Information 3: Species in method comparison**

Table 2. Species used to compare LeafByte, ImageJ, BioLeaf, grid quantification, and visual quantification.

| **Scientific name** | **Common Name** | **Family** | **Code** | **Leaf shape** |
| --- | --- | --- | --- | --- |
| *Asclepias incarnata* | Swamp milkweed | Apocynaceae | AI | lanceolate |
| *Campanula Americana* | American bellflower | Campanulaceae | CA | ovate |
| *Catharanthus roseus* | Madagascar periwinkle | Apocynaceae | MP | lobed |
| *Solidago hybrida 'Dansolitlem'* | Little lemon goldenrod | Asteraceae | SH | lanceolate |
| *Iris ensata* | Japanese iris | Iridaceae | IE | lanceolate |
| *Tropaeolum majus* | Garden nasturtium | Tropaeolaceae | TM | round |
| *Cornus sericea 'Flaviramea'* | Yellow twig dogwood | Cornaceae | CO | ovate |
| *Salvia apiana* | White sage | Lamiaceae | SP | ovate |
| *Primula vulgaris* | Primrose | Primulaceae | PB | ovate |
| *Quercus rubra* | Red oak | Fagaceae | QR | lobed |
| *Thalictrum* 'Elin' | Meadow rue | Ranunculaceae | TH | lobed |
| *Ginkgo biloba* | Ginkgo | Ginkgoaceae | GB | lobed |
| *Sideritis syriaca* | Mountain tea | Lamiaceae | SS | lanceolate |
| *Hydrangea quercifolia* | Oakleaf hydrangea | Cornales | PB | lobed |
